# Mass spectrometrists should search for all peptides, but assess only the ones they care about

**DOI:** 10.1101/094581

**Authors:** Adriaan Sticker, Lennart Martens, Lieven Clement

**Affiliations:** Department of Applied Mathematics, Computer Science & Statistics, Ghent University, Belgium; Medical Biotechnology Center, VIB, Ghent, Belgium; Department of Biochemistry, Ghent University, Ghent, Belgium; Bioinformatics Institute Ghent, Ghent University, Ghent, Belgium

**Author notes:** These authors contributed equally to this work and are joint corresponding author, &.

## Abstract

In shotgun proteomics identified mass spectra that are deemed irrelevant to the scientific hypothesis are often discarded. Noble (2015)^1^ therefore urged researchers to remove irrelevant peptides from the database prior to searching to improve statistical power. We here however, argue that both the classical as well as Noble’s revised method produce suboptimal peptide identifications and have problems in controlling the false discovery rate (FDR). Instead, we show that searching for all expected peptides, and removing irrelevant peptides prior to FDR calculation results in more reliable identifications at controlled FDR level than the classical strategy that discards irrelevant peptides post FDR calculation, or than Noble’s strategy that discards irrelevant peptides prior to searching.

## 1 Introduction

Reliable peptide identification is key to every mass spectrometry-based shotgun proteomics workflow. The growing concern on reproducibility triggered leading journals to require that all peptide-to-spectrum matches (PSMs) are reported along with an estimate of their statistical confidence. The false discovery rate (FDR), i.e. the expected fraction of incorrect identifications, is a very popular statistic for this purpose. In many experiments, however, researchers want to focus on proteins of particular pathways, or few organisms in a metaproteomics sample. Hence, a large fraction of identified peptides are deemed irrelevant for their scientific hypothesis. Considering all PSMs induces an overwhelming multiple testing problem, which leads to few identifications of relevant peptides and thus to underpowered studies.^2^ Currently, there is much debate on the optimal search strategy to boost the statistical power within this context (e.g. Noble, 2015^1^ and http://www.matrixscience.com/nl/201603/newsletter.html).

The common approach, here referred to as the search-all-assess-all (all-all) strategy, involves (1) searching against all expected peptides in a sample, (2) calculating an FDR for each PSM and, (3) filtering irrelevant peptides from the candidate list. Recently, Noble (2015)^1^ pointed out that this strategy is suboptimal because many unnecessary hypotheses are evaluated. Therefore, he proposed to remove irrelevant peptides from the database prior to searching. He argues that a search-subset-assess-subset (sub-sub) strategy improves the statistical power in two ways: (1) each individual spectrum is tested against less candidate sequences and, (2) some spectra that originally matched to proteins that were not of interest will lack a match in the subset search, decreasing the number of PSMs for which an FDR estimate has to be provided.

However, we show that both the all-all and sub-sub strategy are suboptimal and often lead to poorly controlled FDR. From a statistical perspective the all-all strategy FDR is biased as the fraction of incorrect PSMs can differ substantially between the complete set and the subset leading to too conservative or too liberal PSM lists. We also show that the sub-sub strategy forces many good spectra derived from peptides deemed irrelevant to instead match subset peptides. This because the irrelevant peptides were removed from the database prior to searching. Noble correctly pointed out that this issue will not lead to statistical problems as long as a correct FDR procedure is adopted. However, we illustrate that the popular target-decoy FDR procedure^3^ cannot avoid these statistical problems when the sub-sub search strategy is adopted on small to moderate sized subsets. We also argue that the Noble approach still sacrifices statistical power by testing more hypotheses than necessary, i.e. PSMs that would match well to irrelevant peptides in the complete search could actually be discarded because it is highly unlikely that these are subset peptides.

We therefore propose a search-all-assess-subset (all-sub) strategy by (1) searching the mass spectra against a database with all proteins that are expected in the sample, and (2) discarding PSMs matching to irrelevant peptides in the complete search prior to (3) FDR calculation, which has the promise to further boost the statistical power. The filtering strategy in step (2) is independent from the subsequent data analysis steps and can reduce the multiple testing problem considerably without compromising the FDR calculation.^4^

## 2 Case studies

We first evaluate all three strategies on a *Pyrococcus furiosus* dataset with 15,365 high-resolution spectra^5^ (see Supplementary Text). The complete proteome contains 2,051 proteins. We assess 36 subsets ranging from 17 to 381 proteins based on their Gene Ontology (GO) annotations. The spectra are searched with the MS-GF+ search engine^6^ and the FDR is calculated with the target-decoy approach^3^ (TDA, see Supplementary Text). Below, we focus on 175 proteins belonging to the GO term “cytoplasm”. Results for all remaining subsets can be found in supplementary.

In figure 1 we illustrate that the fraction of incorrect target PSMs (*π*_0_) in the complete search is substantially different from the one in the cytoplasm subset. Based on the TDA approach we estimate that 13.9% of the target PSMs are incorrect hits when adopting the all-all search, while the actual fraction in the subset is probably lower, i.e *π*_0_ = 7.2% as estimated with the all-sub strategy. This is also reflected in the distributions of the all-all and the all-sub MS-GF+ scores in figure 1, which are bimodal. The first mode, corresponding to incorrect PSMs, is much higher for the all-all strategy than for the all-sub method. Hence, the FDR cutoff using the all-all search strategy is probably too conservative for the cytoplasm example. This is also reflected by the increased number of subset PSMs that are returned by the all-sub method (2,578 vs 2,553 PSMs). However, the FDR of the all-all method can also be too liberal, i.e. when the fraction of incorrect PSMs in the subset is higher that the one in the complete search (e.g. the ATPase activity subset in supplementary Fig. 10). Hence, the FDR in the all-all strategy is often not representative for that of the subset leading to suboptimal PSM lists, which can be either too long or too short depending on the scenario.

**Figure 1:**
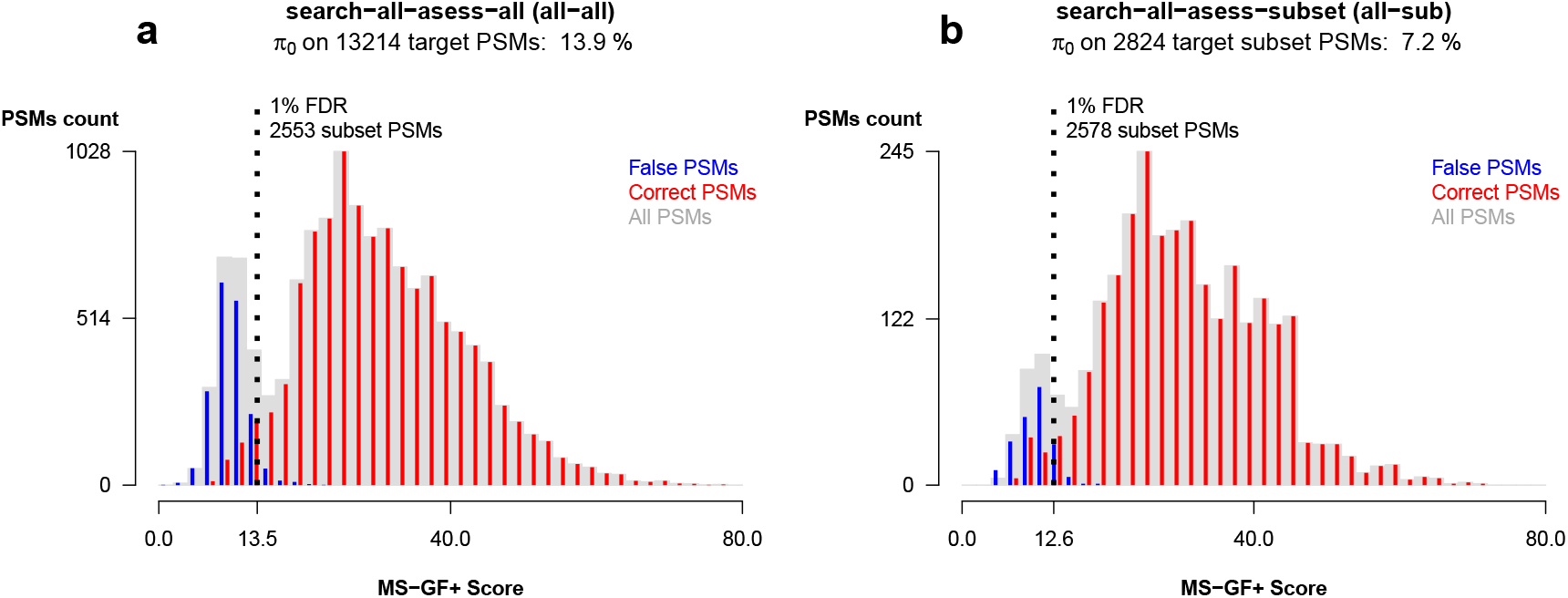
Histograms of MS-GF+ scores (grey) with estimated number of correct PSMs (red, #target - #decoys) and incorrect PSMs (#decoys, blue), 1% FDR cutoff (dashed line). The left modes in the two distributions show that the fraction of null PSMs is higher in the all-all (a) than in the all-sub (b) search suggesting an incorrect FDR control, which is too conservative for the all-all strategy.

Figure 2 illustrates that the Noble sub-sub method is also suboptimal. Noble argues that his strategy leads to a decrease of the number of PSMs that has to be tested. We indeed observe that many target PSMs found in the all-all strategy are no longer matched or matched against a decoy sequence in the sub-sub strategy (left panel of Fig. 2), i.e. only 8,877 target PSMs remain. However, we show that the sub-sub method forces many spectra to match to incorrect sequences, which is increasing the fraction of incorrect PSMs to be tested. In the sub-sub strategy 5,831 out of 8,877 target PSMs (*π*_0_ = 65.7%, Fig 2. a) are expected to be incorrect compared to only 1,839 out of 13,214 PSMs (*π*_0_ =13.9%, Fig. 1 a) in the all-all strategy. Because of the higher fraction of incorrect PSMs in the sub-sub strategy, the score cutoff at 1% FDR increases from 13.5 to 15.0. A similar increase of the score cutoff at 1% FDR was also observed in all other GO subsets (see Supplementary Fig. 1-35). Due to this higher score cutoff, the sub-sub method will return a lower number of subset PSMs that match to the same peptide as in the all-all search strategy. Despite the more stringent cutoff, however, the sub-sub strategy returns more subset PSMs in 13 out of the 36 GO subsets. This happens because several spectra, derived from a peptide that was deemed irrelevant are now forced to match subset peptides. In the cytoplasm example (Fig. 2. b), the sub-sub PSM list contains many such PSMs that switched peptide, i.e. 53 on 2,559 PSMs at 1% FDR. This actually amounts to 2.2% of all returned PSMs, which strongly suggests an error rate of at least twice the adopted 1% FDR. Indeed, most of these switched PSMs are likely to be false positives: their MS-GF+ score is always lower for the sub-sub match than for the all-all match (see Fig. 2 b). The same behavior is also observed for many of the other subset searches, and this both at 1% and 5% FDR (Supplementary Fig. 36). Hence, the increased number of PSMs returned by the sub-sub method are highly questionable at best. We hypothesize that this is partially induced by an erratic behavior of TDA when dealing with small search libraries. This erratic behavior is illustrated in figure 2 a) where an unexpected enrichment in the number of expected correct PSMs can be observed at low MS-GF+ scores. This suggests that low quality spectra in a restricted search space tend to match incorrectly to target sequences because there is insufficient sequence variation in the decoys. This in turn leads to a poorly controlled FDR. The latter concern was also mentioned by Noble^1^ and considerably reduces the reliability of the sub-sub search method for small to moderate subsets. In contrast, our proposed all-sub strategy (Fig. 1. b) also reduces the multiple testing problem considerably, but does not force good spectra to incorrectly match with subset peptide sequences because we search against all possible sequences.

**Figure 2:**
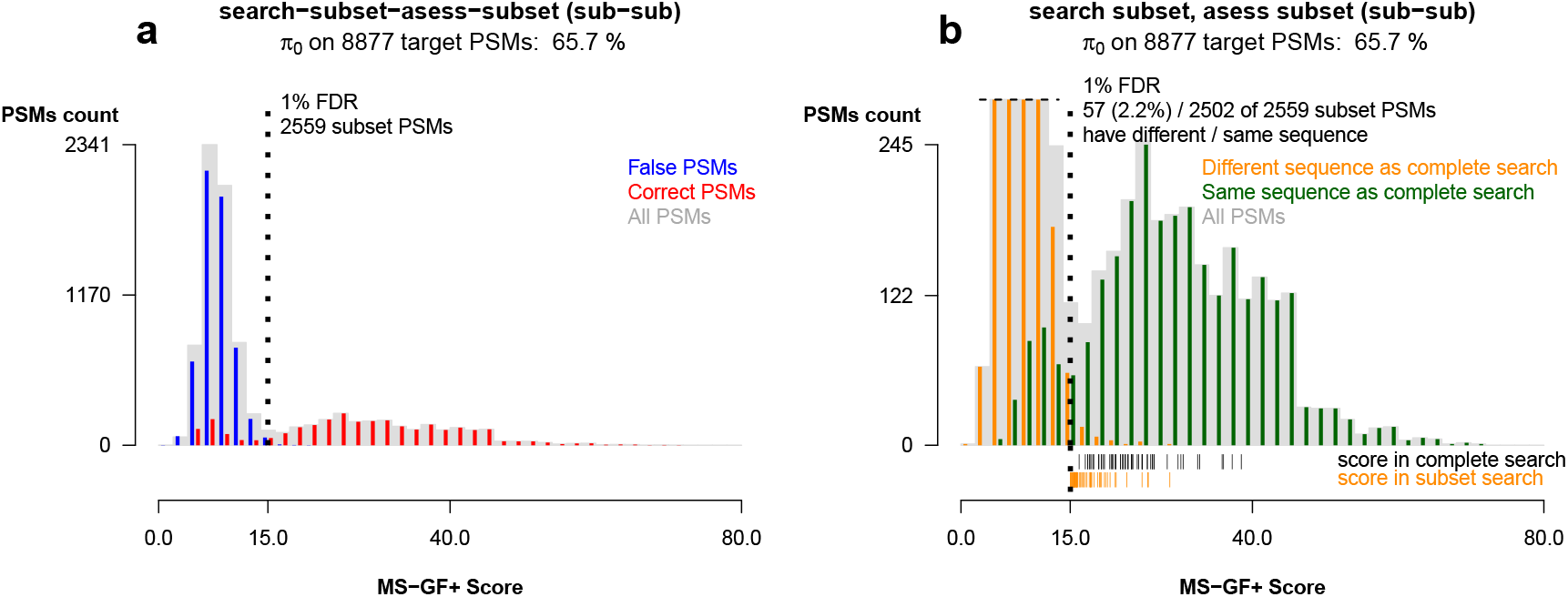
Histograms of MS-GF+ scores (grey) for the Noble search-subset-assess-subset strategy. a) estimated number of correct PSMs (red, #target - #decoys) and incorrect PSMs (#decoys, blue). Compared to all-all strategy in figure 1 a) the score cutoff at 1% FDR (vertical dashed line) shifts to higher values, but still more subset PSMs are found. It also shows that many PSMs are forced on incorrect subset PSMs (huge first mode of the distribution). b) common PSMs (green) and PSMs that switched peptides sequences (orange) in the sub-sub strategy as compared to the all-all strategy in figure 1 a). These switched PSMs all have lower scores than in the all-all search and are therefore questionable at best (black and orange rug plot below histogram).

In a second example we evaluated the methods on a malaria parasite *Plasmodium Falciparum* dataset containing 55,036 spectra.^7^ The *Plasmodium Falciparum* proteins in the dataset were captured from a mixed culture with human red blood cells. Hence, the samples are likely to be contaminated with a considerable amount of human proteins. We assessed two subsets of interest:the plasmodium subset and the human subset consisting of 5,136 and 20,201 proteins, respectively. We do not expect problems with the TDA-FDR estimation in the sub-sub strategy because each subset contains more than 1,000 proteins.^1^ This is confirmed in Supplementary Figures 37 and 38, which do not show bimodal patterns in the distribution of the expected correct PSMs. Similar to the *Plasmodium* example in the paper of Noble,^1^ the sub-sub approach retrieves more PSMs than the all-all strategy at 1% FDR (14,277 vs 1,3869 PSMs, respectively). However, the opposite is observed for the human subsample (6,108 vs 6,474 PSMs).

Again, the all-all FDR-control seems inadequate. in the complete set and the subset seems to differ considerably, i.e. the all-all strategy seems too conservative in the plasmodium subset (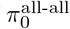 and 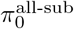 = 24.5%, Supplementary Fig. 38), and too liberal in the human subset (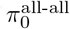 = 38.1% and 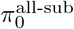 = 54.6%, Supplementary Fig. 39).

The Noble sub-sub strategy, on the other hand, is more conservative than our all-sub strategy at 1% FDR and returned less subset PSMs for both subsets (Supplementary Fig. 38-39). Moreover, a considerable fraction of the returned PSMs still switched peptides between the all-all and sub-sub searches, i.e. 0.6% in the *Plasmodium* and 1.6% in the human subset at 1% FDR. As before, these switched PSMs are likely to contain many induced false positives. In the *Pyrococcus* example, we observed that the higher *π*_0_ of the sub-sub strategy increased the score cutoffs at 1% FDR. Here, we observe a trade-off between the change in *π*_0_ and in the decoy distribution (Supplementary Fig. 38, panel a vs b, blue bars). The latter might be due to differences in overall protein composition in *Pyrococcus* and human subsets. In the *Plasmodium* subset the FDR cutoff still decreases from 20.5 to 20.2 despite the increase in *π*_0_ (from 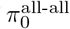 = 38.1% to 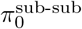 = 51.1%). This explains the increase in the number of returned subset PSMs as compared to the all-all method, i.e. an additional 324 subset PSMs with the same label are returned but they come at the expense of 84 switched PSMs. Supplementary Figure 39 (panel a vs b) shows a much larger *π*_0_ increase for the human subset (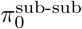 = 72.0%). This cannot be compensated by distributional changes of the null component, resulting in a higher threshold than with the strategies based on the complete search. The higher threshold leads to a lower number of returned human PSMs in common with the all-all search, and a high fraction of switched PSMs (1.6% at the 1% FDR level). Hence, even for large subsets the sub-sub TDA FDR seems questionable.

## 3 Challenges and future directions

Both the *Pyrococcus* and *Plasmodium* example clearly indicate that mass spectrometrists will benefit from searching for all peptides, but by only assessing the ones they care about. Our new strategy returns more PSMs than the Noble approach and can be expected to provide a better FDR control within peptide subsets than the two leading strategies, which discard peptides prior to searching or post FDR calculation. Adopting the target decoy approach in our all-sub strategy involving small subsets, however, often provides unstable FDR estimates due to a considerable sample to sample variability in the number of subset decoys above a particular score cutoff. For instance, the transmembrane transport GO subset has few PSMs and only 7 decoy hits. The subset is higher than in the complete search and a more stringent MS-GF+ score cutoff seems to be required. But due to the specific emperical distribution of the decoy scores the subset TDA still results in a lower cutoff than in the all-all approach (Supplementary Fig. 27).

We observe that the location and shape of (1) the decoy distribution and (2) the estimated correct distributional component of the target distribution in the all-all and all-sub strategy are very similar (e.g. Supplementary Fig. 1-35, 38-39), indicating that the overall properties of the PSMs in subsets and the complete set remain alike. We therefore propose to exploit the full information available in the complete search for improving the estimation of the all-sub FDR.

Hence, we can estimate the distributional components using the complete search and only have to rely on the subset decoy and target PSMs for the calculation of (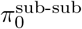). The latter FDR estimate is more stable as it involves all subset decoys in contrast to the all-sub TDA, which only consists of the relatively few subset decoys above a score cutoff. The reweighted FDR indeed results in a more stringent MS-GF+ score cutoff for the transmembrane transport GO subset (Supplementary Table 1).

We also developed a user-friendly web-based tool in R^8^ that provides (1) the all-sub FDR, (2) the rescaled all-all FDR and (3) diagnostic plots for assessing the location-scale assumption. (http://iomics.ugent.be/saas/ and Supplementary Code) In our application, *π*_0_ is estimated based on the ratio of the number of subset decoys and the number of subset targets in a concatenated target-decoy search. We feel that our approach can be further optimized for small subsets by using the location and shape assumption explicitely when estimating *π*_0_.

